# ATP-release drives inflammation with lysophosphatidylcholine

**DOI:** 10.1101/2020.12.25.424382

**Authors:** Sana Ismaeel, Ayub Qadri

## Abstract

Lysophosphatidylcholine (LPC), a dominant lipid component of oxidized low-density lipoprotein, plays a major role in inflammation associated with atherosclerosis and neurodegenerative disorders. It activates inflammatory responses from macrophages, neuronal cells and endothelial cells. However, the exact mechanism by which LPC promotes inflammation remains incompletely understood. Here, we show that the production of inflammatory cytokines and cytotoxicity with LPC are both critically dependent on its ability to bring about release of ATP from cells. The induction of caspase-1-mediated IL-1β-release with LPC from TLR-primed macrophages and neuronal cells is reduced in presence of ATP-hydrolyzing enzyme, apyrase and the inhibitors of purinergic signaling. ATP released from LPC-treated cells also promotes an IL-12p70^hi^, low phagocytic and poorly co-stimulatory phenotype in macrophages in a caspase-1 – independent manner. Treatment with apyrase reduces production of inflammatory cytokines with LPC *in vivo*. These findings reveal a previously unappreciated pathway for generation of inflammatory responses with LPC, and these have significant implications for therapeutic intervention in chronic inflammatory disorders promoted by this lipid.

## Introduction

LPC, an important constituent of normal plasma, is produced by the action of phospholipase A2 on phosphatidylcholine that is the major lipid in the cell membrane (*1*). It performs a number of physiological functions in different cell types by engaging the G-protein coupled receptors, G2A and GPR4, with the former being more extensively studied (*2*, *3*). LPC also forms a significant component of the oxidized low-density lipoprotein (Ox-LDL) and is considered to be a critical player in the atherogenic activity of this lipoprotein (*4*). It is present in abundance in platelet-derived microvesicles that cause vascular inflammation (*5*). Increased PLA2 activities have been associated with acute and chronic inflammation (*6*), and high LPC concentrations have been shown to bring about demyelination in various models of neurodegeneration (*7*). *In vitro*, LPC produces inflammatory responses from macrophages, neuronal cells and endothelial cells, and it also bring about cellular cytotoxicity (*8*–*10*). Recent studies have suggested that a major pathway by which this lipid can contribute to chronic inflammation is through activation of caspase-1 – mediated release of IL-1β *via* stimulation of the Nlrp3 and Nlrc4 inflammasomes (*8*, *9*). Its activation of Nlrp3 has also been implicated in the promotion of foamy macrophages, which are known to be abundant in atherosclerotic plaques (*10*). Due to its association with chronic inflammatory disorders, LPC is now recognized as one of the major damage associated molecular patterns (DAMP) (*11*). However, the exact mechanism by which this lipid promotes inflammatory responses during different disease states remains incompletely understood.

In the present study, we demonstrate that LPC triggers release of another DAMP, ATP, from macrophages and neuronal cells, and reduces the threshold of one or more P2X receptors to this DAMP to bring about cell death and release of caspase-1-dependent IL-1β. This extracellular ATP (eATP) also promotes production of IL-12p70 from TLR-sensitized macrophages, reduces phagocytic ability of macrophages and surface expression of MHC class II, CD 86 and several other molecules on these cells. All these responses are abrogated *in vitro* as well as *in vivo* in presence of ATP-hydrolyzing enzyme, apyrase. These results suggest that ATP-release is the crucial first step in the induction of cell death and generation of inflammatory responses with LPC.

## Results

### LPC induces release of ATP to bring about production of IL-1β

To understand the mechanism by which LPC activates production of caspase-1 – dependent IL-1β, we investigated the role of GPCRs, which are known to regulate a large number of cellular responses with this lipid (*12*), in this activation. TLR – primed macrophages were stimulated with LPC in presence of GPCR inhibitors, pertussis toxin (PT) and suramin, and IL-1β was analyzed in cell culture supernatants. LPC activated release of IL-1β from LPS-primed mouse peritoneal macrophages, human monocytic cell line, THP-1 and murine microglial cell line, N-9, in a dose-dependent manner (Fig. 1 A; Supplementary Fig. 1 A, D). This response was not seen with phosphatidylcholine (PC; Fig. 1 B). The release of IL-1β with LPC was not inhibited in presence of PT suggesting that activation of caspase-1 with this lipid might be independent of Gαi (Fig. 1 C; Supplementary Fig. 1 B, E) (*13*, *14*). On the other hand, treatment with suramin inhibited IL-1β release from LPC-stimulated cells (Fig. 1 D; Supplementary Fig. 1 C, F). As suramin is also known to inhibit signaling through purinergic receptors, and LPC had been previously shown to sensitize cells to ATP (*15*, *16*), we considered the possibility that LPC might activate production of IL-1β by bringing about release of ATP from cells and simultaneously priming cells to respond to this DAMP. To test this possibility, we carried out LPC-stimulations in presence of ATP - hydrolyzing enzyme, apyrase. This enzyme reduced LPC-stimulated IL-1β secretion from peritoneal mouse macrophages, THP-1 cells and N-9 cells in a dose-dependent fashion (Fig. 1 E, F; Supplementary Fig. 2). As expected, apyrase also inhibited IL-1β release from TLR-primed cells stimulated with high concentrations of ATP (Fig. 1 G). However, it did not reduce IL-1β production from cells activated with nigericin, infected with *Salmonella* or transfected with flagellin protein (Fig. 1 G). These results suggested that the induction of ATP-release might be critical for generation of IL-1β with LPC.

**Figure 1:**
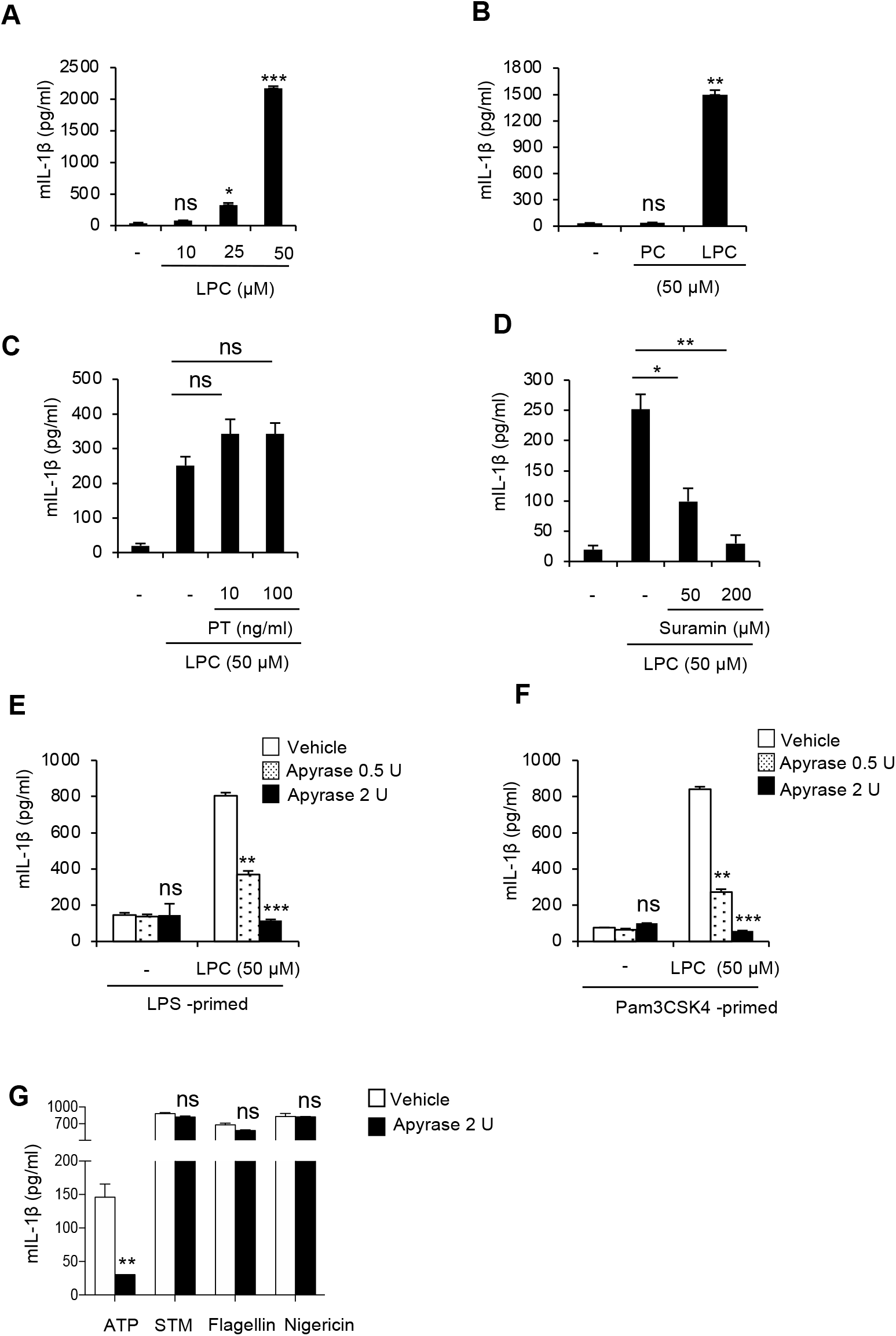
LPC – mediated caspase-1 – dependent IL-1β production is abrogated in presence of apyrase. (**A, B**) Mouse peritoneal macrophages plated at a density of 5×10^5^ cells per well in RPMI −1640 supplemented with 10% FCS (RPMI-10) were stimulated with LPS (1 μg / ml) for 6 h. Cells were washed with serum-free RPMI-1640 (RPMI) and incubated with different concentrations of LPC or PC diluted in RPMI. Cell-free supernatants were collected after 3 h and IL-1β was determined by ELISA. (**C, D**) LPS-primed mouse macrophages were pretreated for 30 minutes with pertussis toxin or suramin before stimulating with LPC (50 μM) for 3 h at 37°C. Cell - free supernatants were analyzed for IL-1β. (**E, F**) Cells were primed with LPS or Pam3CSK4 (1μg / ml) for 6 h in RPMI-10, washed and incubated with apyrase for 30 minutes followed by LPC (50 μM) for 3 h. IL-1β was determined in cell supernatants by ELISA. (**G**) Cells were activated with LPS, washed and treated with apyrase for 30 minutes before stimulating with ATP, *Salmonella* Typhimurium (STM), Nigericin or transfected with *Salmonella* Typhi flagellin. Cell-free supernatants were collected after 1 h (for infection and nigericin) or 3 h (for ATP and flagellin), and IL-1β was determined by ELISA. Data are representative of 2-3 independent experiments. *p< 0.05; ** p<0.01; *** p<0.001; ns – not significant.

Analysis of ATP in the extracellular medium showed that cells treated with LPC but not PC triggered release of this endogenous DAMP in a dose-dependent manner, with levels peaking at about 10 minutes (Fig. 2 A, B). The amount of ATP that was released by LPC-treated cells was in low nanomolar range, which is much lower than the millimolar concentration of ATP that normally activates caspase-1 from TLR-primed cells (Fig. 2 A, C) (*17*). Therefore, in addition to bringing about release of ATP, LPC also primed cells to respond to low levels of eATP. Analysis of different derivatives of LPC showed that longer chain saturated derivatives triggered more ATP-release from cells and brought about increased production of IL-1β from TLR-primed cells (Fig. 2 D, E). None of the derivatives however elicited IL-1β in unprimed cells (data not shown). ATP-release, as reported previously, was also seen in cells infected with *Salmonella* but treatment with apyrase did not inhibit release of IL-1β from these cells indicating that eATP did not participate in generation of this cytokine with this pathogen (Fig. 2 F; Fig. 1 G) (*18*, *19*).

**Figure 2:**
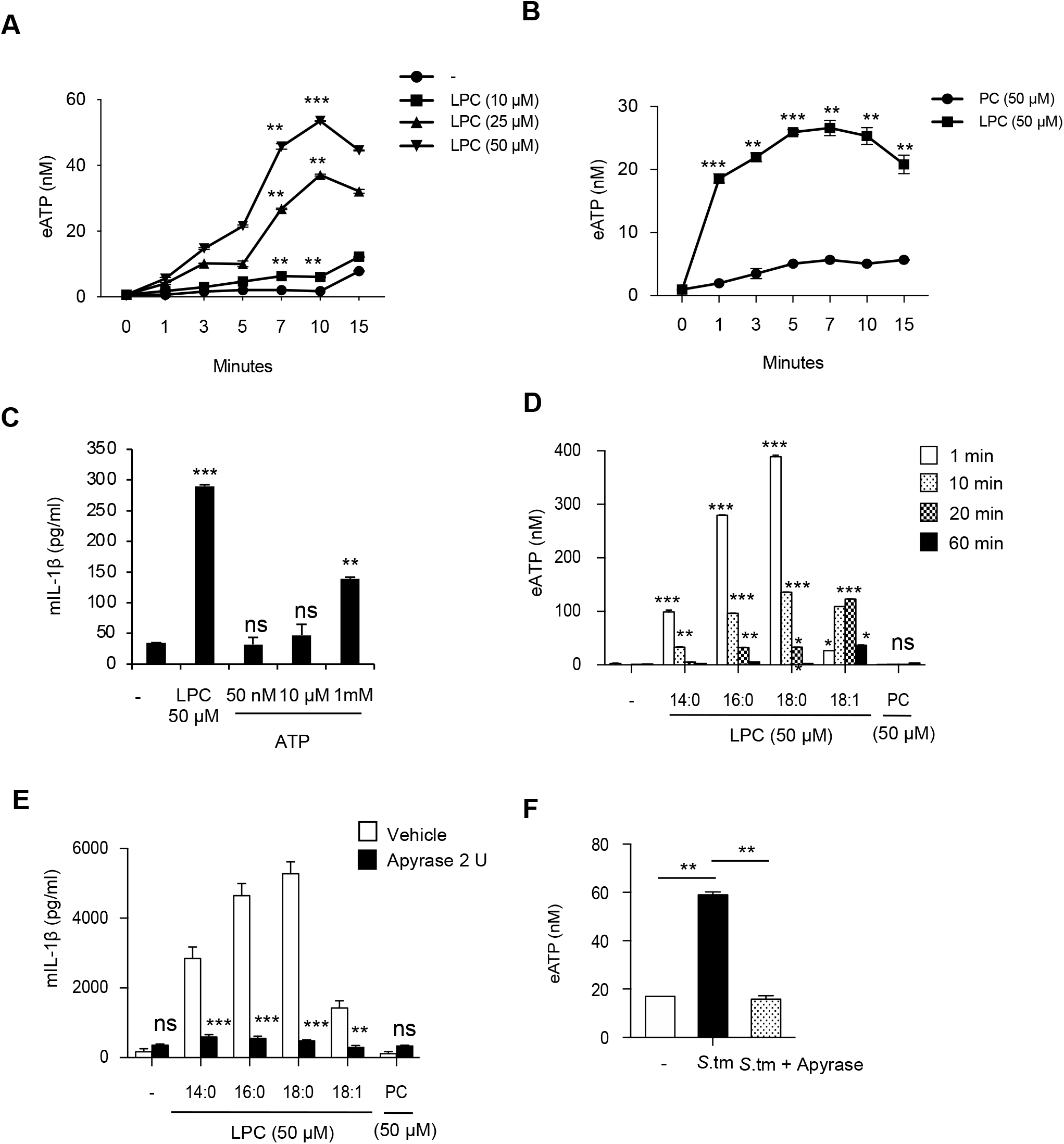
LPC activates release of ATP for IL-1β production. (**A, B**) Mouse peritoneal macrophages plated at 3×10^5^ cells per well in RPMI-10 were washed free of serum and stimulated with LPC or PC in RPMI. The supernatants were collected at different time points and boiled at 90°C for 2 minutes. ATP was quantified using ATP-bioluminescence assay kit. (**C**) LPS - primed mouse macrophages were stimulated with LPC (50 μM) or different concentrations of ATP. After 3 h, cell-free supernatants were analyzed for IL-1β by ELISA. (**D**) Mouse macrophages were washed free of serum and stimulated with different derivatives of LPC in RPMI. The supernatants were collected at different time points and ATP was quantified using ATP-bioluminescence assay kit. (**E**) LPS - primed mouse macrophages were pretreated with apyrase before stimulating with PC or different derivatives of LPC (50 μM). After 3 h, cell-free supernatants were analyzed for IL-1β by ELISA. (**F**) Mouse peritoneal macrophages were infected with *Salmonella* Typhimurium (STM) in the absence or presence of apyrase for 1 h and ATP was measured in the supernatants. Data are representative of 2-3 independent experiments. *p< 0.05; ** p<0.01; *** p<0.001; ns – not significant.

Consistent with a recent study, the release of IL-1β in response to stimulation of cells with LPC was also reduced in presence of intracellular calcium chelator BAPTA-AM, and the inhibitors of Syk, potassium efflux and JNK (Supplementary Fig. 3) (*8*).The inhibition with BAPTA-AM prompted us to look at the role of eATP in mobilizing intracellular calcium. Treatment of mouse macrophages with LPC but not PC activated release of calcium which was abrogated in presence of apyrase suggesting that eATP was solely responsible for bringing about calcium mobilization in LPC-stimulated cells (Supplementary Fig. 4)

### ATP-release is essential for production of IL-1β with LPC *in vivo*

Intraperitoneal administration of LPC to mice did not show any detectable increase in the basal levels of IL-1β in the peritoneal lavage (Fig. 3 A). However, sensitization of mice with LPS before administering LPC resulted in significant enhancement in the levels of IL-1β (Fig. 3 A). This increase was reduced in mice which received apyrase along with LPC in the peritoneal fluid as well as serum, establishing a crucial role for eATP in the induction of IL-1β with this lipid *in vivo* (Fig. 3 B, C). At the concentration and time used in this study, LPS on its own did not increase IL-1β above background level in the peritoneum.

**Figure 3:**
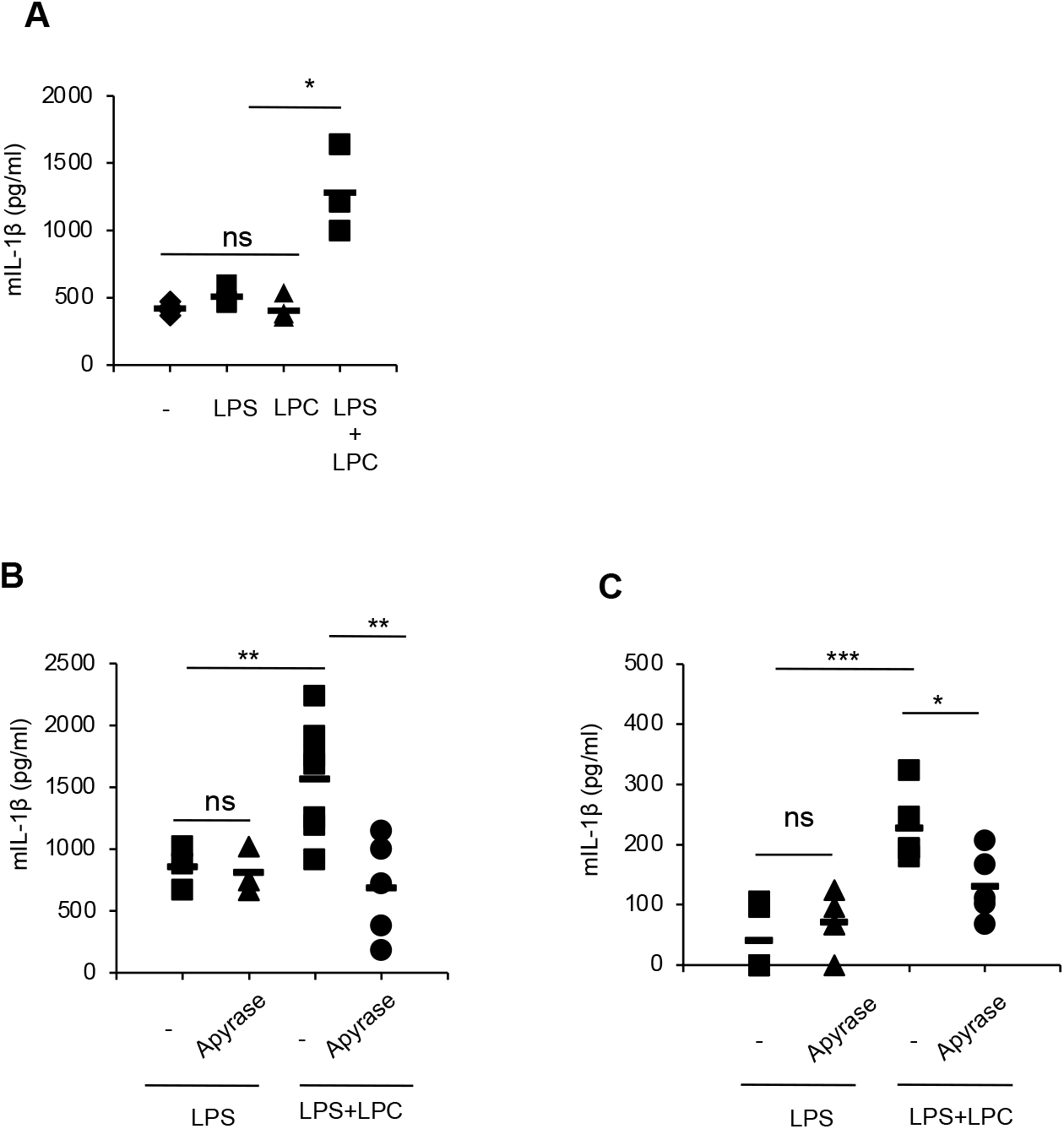
LPC causes ATP-release to produce IL-1β *in vivo*. (**A**) 6-8-week-oldC57BL/6 mice were injected intraperitoneally with LPS (7μg per mouse). After 10 h, LPC (200 μg) dissolved in RPMI was injected in the absence or presence of apyrase (15 U). Peritoneal lavages (**A, B**) and sera (**C**) were collected after 2 h and 4 h respectively; and the levels of IL-1β were determined by ELISA. Data are representative of 2-3 independent experiments. *p< 0.05; ** p<0.01; *** p<0.001; ns – not significant.

### Pannexin-1 and P2XR are required for LPC-mediated release of ATP and IL-1β

The results described above suggested that LPC triggers release of cellular ATP which in turn brings about release of IL-1β very likely through stimulation of purinergic signaling. A major mechanism by which ATP is released from cells is through activation of the pannexin-1 channel (*20*–*22*). We analyzed possible role of this channel by determining release of ATP and IL-1β from LPC-stimulated cells in presence of the inhibitors of this channel. Treatment of cells with pannexin-1 inhibitors, probenecid and carbenoxolone, before activating with LPC reduced release of ATP indicating that this release was brought about through opening of this channel (Fig. 4 A, B). Importantly, pannexin-1 inhibitors also reduced release of IL-1β from cells activated with LPC (Fig. 4 C, D).

**Figure 4:**
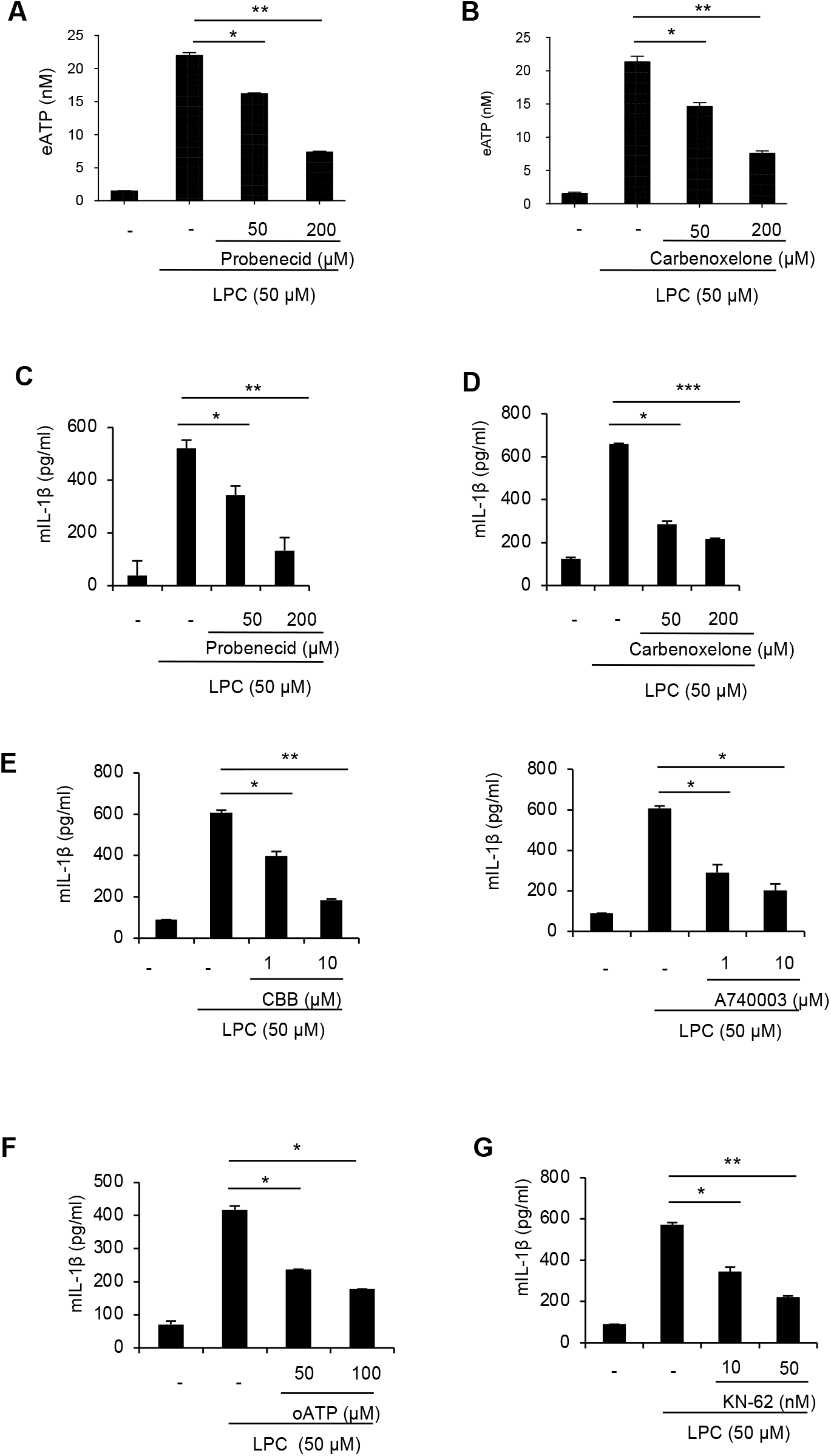
Pannexin-1 and P2XR are essential for generation of IL-1β from macrophages stimulated with LPC. (**A, B**) Murine macrophages were washed free of serum and treated with pannexin-1 inhibitors, probenecid and carbenoxolone, for 1 h. Cells were then stimulated with LPC for 5 min, supernatants were collected and boiled at 90°C for 2 minutes. ATP was quantified using ATP-bioluminescence assay kit. (**C, D**) Murine macrophages were primed with LPS (1 μg/ml) for 6 h in RPMI-10, washed and incubated with pannexin-1 inhibitors, probenecid or carbenoxolone, for 1 h followed by activation with LPC for 3 h. The supernatants were analyzed for IL-1β by ELISA. (**E, F, G**) Cells were primed with LPS and treated with P2XR inhibitors, CBB-G250, A740003 or KN-62 for 30 min or with oATP for 2 h followed by incubation with LPC for 3 h. IL-1β was determined in the cell supernatants by ELISA. Data are representative of 2-3 independent experiments. *p< 0.05; ** p<0.01; *** p<0.001; ns – not significant.

Activation of P2X7 receptor brings about pore formation for permeation of dyes like YO-PRO1 (*23*). LPC caused YO-PRO1 uptake which was abrogated in the presence of apyrase suggesting involvement of P2X7R in LPC-mediated IL-1β release (Supplementary Fig. 5). Consistent with this observation, the production of IL-1β from TLR-primed cells stimulated with LPC was reduced in presence of inhibitors of purinergic signalling, oxidized ATP (a P2XR inhibitor) or CBB-G250, A740003, and KN-62 (specific P2X7R inhibitors) (Fig. 4 E, F, G; Supplementary Fig. 6). These results clearly established a role for one or more purinergic receptors in bringing about release of IL-1β from cells activated with LPC.

### Cell death with LPC is largely caspase-1 independent

LPC has been previously shown to activate caspase-1 through stimulation of Nlrp3 as well as Nlrc4 (*8*, *9*). Nlrp3 is normally expressed at low levels in cells and it is induced following TLR-priming, while Nlrc4 is constitutively expressed (*24*). We asked if LPC could activate cell death in the absence of TLR-priming. Treatment of mouse peritoneal macrophages with LPC resulted in release of LDH in a dose - dependent manner (Fig. 5 A). TLR-priming of cells did not significantly increase this release with LPC (Fig. 5 B). Moreover, neither release of LDH nor that of ATP was inhibited with caspase-1, caspase-3 or pan-caspase inhibitor (Fig. 5 C, D). On the other hand, apyrase completely abrogated release of LDH in response to LPC suggesting that LPC-induced cytotoxicity in macrophages may be driven by eATP independent of caspase activation. (Fig. 5 E). This cytotoxicity was partially reduced in presence of calcium chelators or an inhibitor of purinergic signaling (Fig 5 F). Consistent with its inability to activate release of ATP, PC did not bring about LDH - release above background levels from any of the cell types (data not shown).

**Figure 5:**
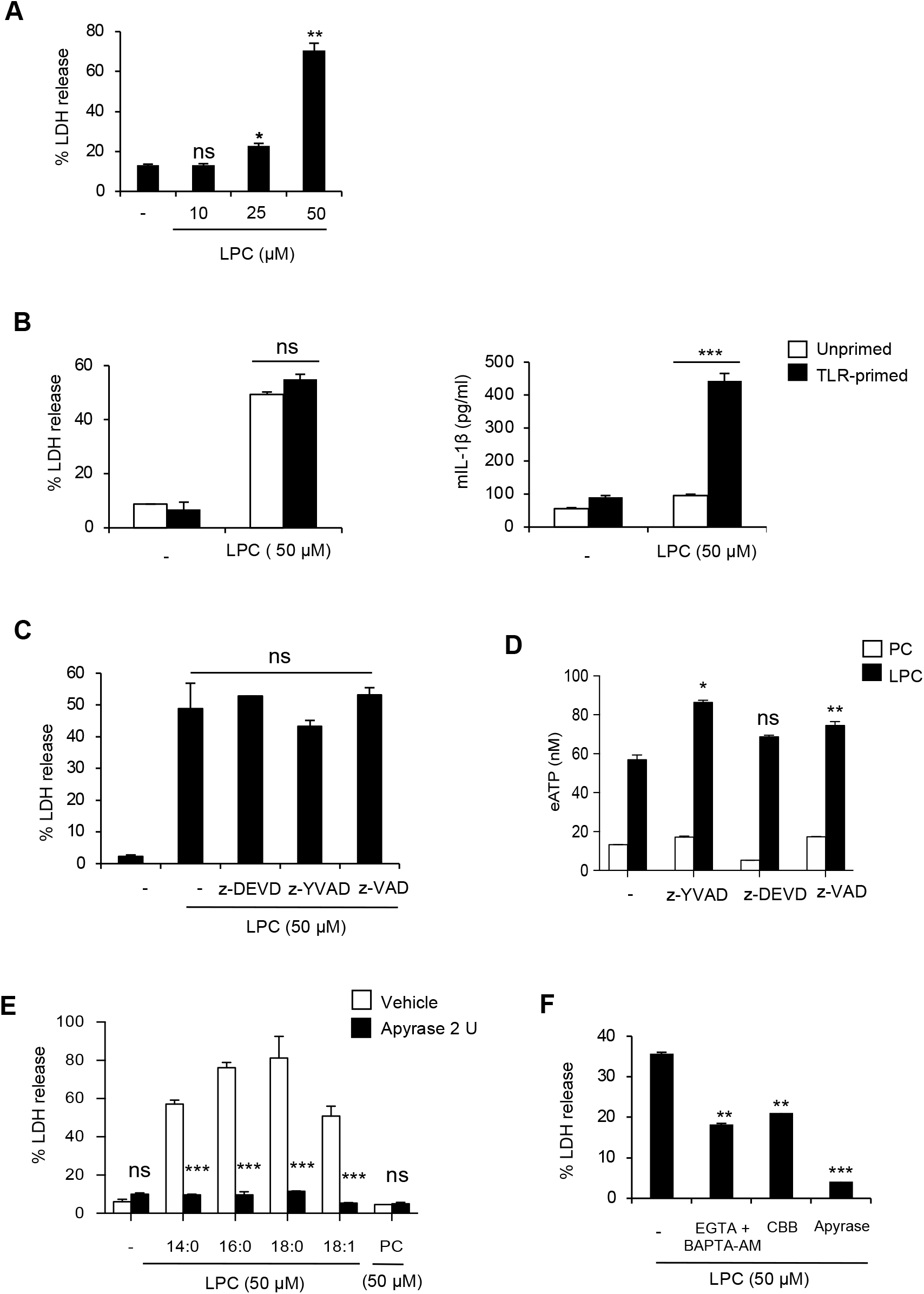
LPC induced ATP causes cell death in a caspase-1 independent manner. (**A**) Mouse peritoneal macrophages were washed with RPMI and incubated with different concentrations of LPC for 3 h. LDH levels were determined in the cell supernatants by LDH-cytotoxicity kit. (**B**) LPS-primed and unprimed macrophages were stimulated with LPC and levels of IL-1β and LDH were determined. (**C**) Mouse peritoneal macrophages were washed with RPMI and pretreated with caspase inhibitors before incubating with LPC for 3 h. LDH levels were determined in the cell supernatants by LDH-cytotoxicity kit. (**D**) Mouse peritoneal macrophages were washed with RPMI and pretreated with caspase inhibitors (50 μM) before incubating with LPC. Supernatant was collected after 5 minutes and boiled at 90°C for 2 minutes. ATP - release was quantified using ATP-bioluminescence assay kit. (**E**) Mouse peritoneal macrophages were washed with RPMI and pretreated with apyrase before incubating with different derivatives of LPC for 3 h. LDH levels were determined in the cell supernatants by LDH-cytotoxicity kit. (**F**) Mouse macrophages were pretreated with EGTA and BAPTA-AM, CBB-G250 or apyrase before stimulating with LPC in serum-free RPMI. LDH levels were determined in the cell supernatants. Data are representative of 3 independent experiments. *p< 0.05; ** p<0.01; *** p<0.001; ns – not significant.

### LPC-triggered ATP promotes an IL-12p70^hi^, low phagocytic and poorly costimulatory phenotype in LPS - primed mouse monocytes

In the course of analyzing caspase-1 – dependent IL-1β release with LPC, it was observed that treatment of TLR-primed but not unprimed peritoneal macrophages with this lipid results in generation of a population of cells with a morphology which resembled that of a mature dendritic cell (DC) (Fig. 6 A). Macrophages primed through TLR4 or TLR2 and stimulated with LPC showed this kind of a phenotype with cells enriched in dendrites (Fig. 6 B). This phenotype was observed as early as 30 minutes after incubation with LPC (data not shown) and at 3 h prominent dendrites could be readily seen. These cells did not take up propidium iodide or Sytox-green indicating that these were resistant to LPC-driven cell death (Fig. 6 C, D). PC did not produce this phenotype (Fig. 6 E). Moreover, LPC did not bring about this transformation in unprimed monocytes (Fig. 6 A). In unprimed cells, treatment with LPC resulted in rounding off of monocytes and prolonged incubation with LPC resulted in detachment of cells from the substratum (Fig. 6 A).

**Figure 6:**
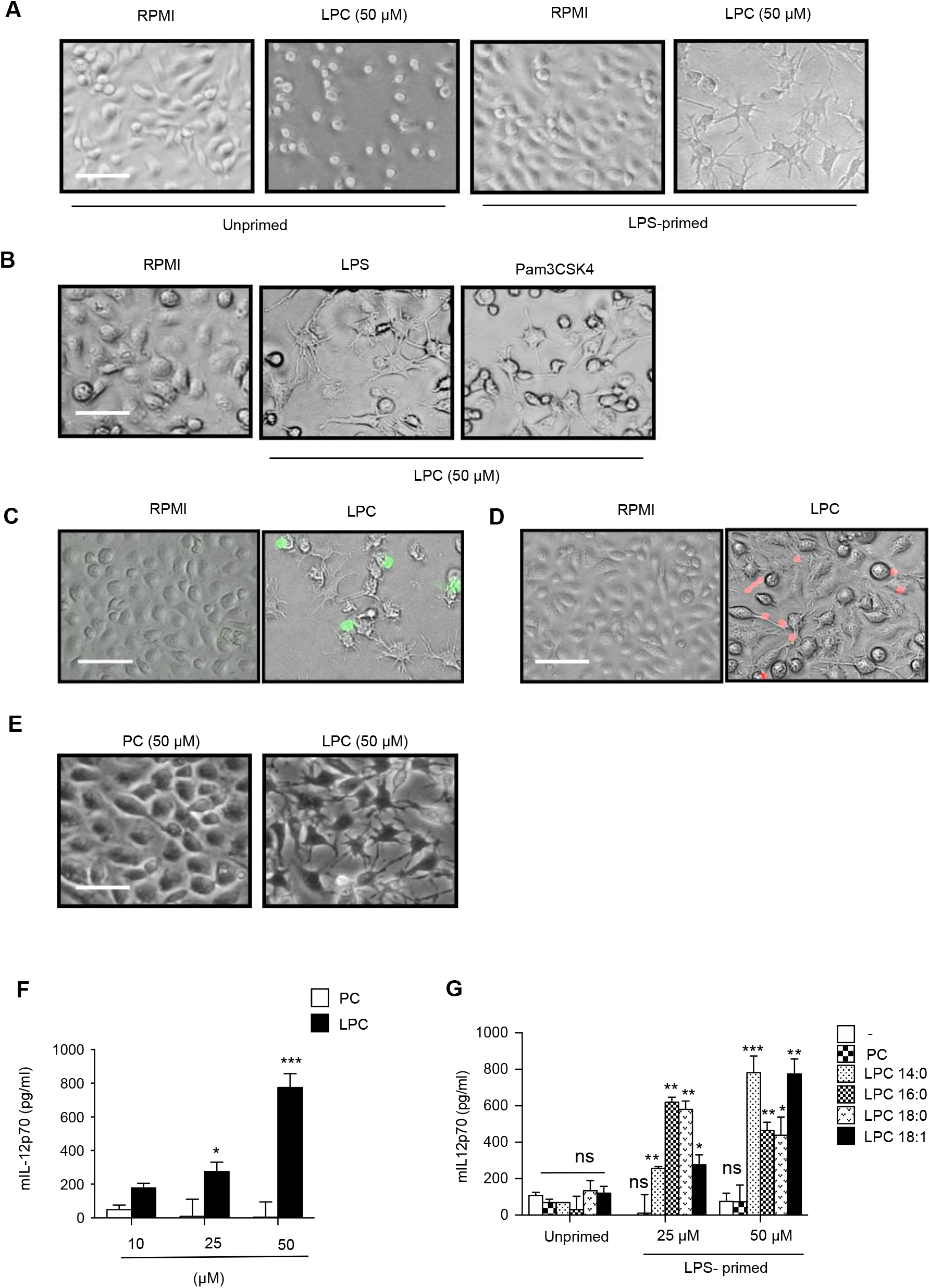
LPC but not PC produces a mature DC-like phenotype and release of IL-12p70 in TLR-primed macrophages. (**A, B, C, D, E**) Mouse peritoneal macrophages plated at a density of 5×10^5^ cells per well in RPMI-10 were stimulated with LPS (1 μg/ml) or Pam3CSK4 (1 μg/ml) for 6 h. Cells were washed with serum-free RPMI and incubated with LPC or PC diluted in RPMI. 3 h later, cells were stained with (**C**) Sytox-green or (**D**) propidium iodide for 15 minutes. Images were captured in a fluorescence microscope (Nikon TE-2000) using ACT software and imported into adobe photoshop. Scale bar-50 μm. (**F, G**) LPS-primed or unprimed mouse macrophages were stimulated with different concentrations of PC or various derivatives of LPC for 3 h at 37°C. Cell - free supernatants were analyzed for IL-12p70. Data are representative of 3 independent experiments. *p< 0.05; ** p<0.01; *** p<0.001; ns – not significant.

The ‘mature dendritic cell like phenotype’ of these macrophages prompted us to analyze secretion of IL-12p70, expression of MHC class II and co-stimulatory molecules like CD80, CD86, CD40 and the ability of these cells to phagocytose particulate material. LPC but not PC increased secretion of IL-12p70 from TLR-primed cells in a dose-dependent fashion (Fig. 6 F). This ability was seen with different derivatives of LPC (Fig. 6 G). Unprimed cells did not secrete any IL-12p70 upon treatment with LPC (Fig. 6 G). LPC 16:0 and LPC 18:0 produced more cell death that is why IL-12p70 secretion was reduced with these two derivatives at higher concentration (Fig. 6 G, Fig. 5 E). Interestingly, while treatment of LPS-primed LPC-treated cells with caspase-1 inhibitor, as expected, reduced IL-1β, it did not inhibit secretion of IL-12p70 (Fig. 7 A). Significantly, however, treatment with apyrase before stimulation with LPC completely abrogated production of IL-12p70 suggesting that eATP produced by LPC-treated cells brought about secretion of this cytokine independent of caspase-1 activation (Fig. 7 B). Similarly, treatment with apyrase but not caspase-1 inhibitor prevented the dendrite^hi^ macrophage phenotype produced by LPC in TLR-primed macrophages (Fig. 7 C).

**Figure 7:**
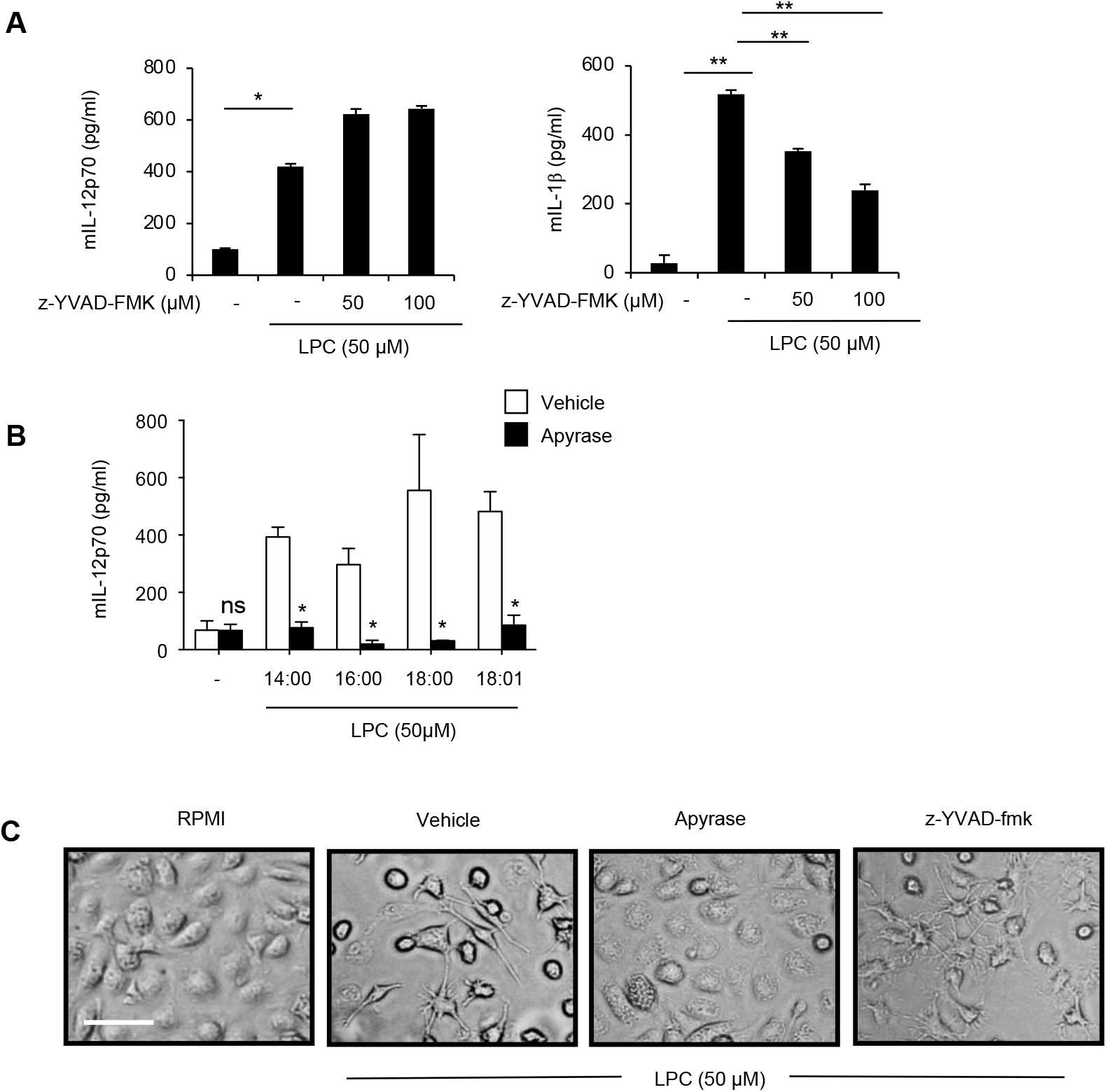
Treatment with apyrase but not caspase-1 inhibitor reduces secretion of IL-12p70 and mature DC-like phenotype. (**A**) LPS-primed mouse peritoneal macrophages were washed with RPMI and pretreated with z-YVAD-FMK before incubating with LPC for 3 h. IL-1β and IL-12p70 levels were determined in the cell supernatants by ELISA. (**B**) LPS-primed mouse peritoneal macrophages were washed with RPMI and pretreated with apyrase before incubating with LPC or its different derivatives for 3 h. IL-12p70 levels were determined in the cell supernatants by ELISA. (**C**) LPS-primed mouse peritoneal macrophages were washed with RPMI and pretreated with apyrase or z-YVAD-fmk before incubating with LPC for 3 h. Images were captured in a fluorescence microscope (Nikon TE-2000) using ACT software and imported into adobe photoshop. Scale bar-50 μm. Data are representative of 2-3 independent experiments. *p< 0.05; ** p<0.01.

To further characterize these cells, we also analyzed surface expression of MHC class II and co-stimulatory molecules. As expected, LPS showed induction of MHC class II, CD86, CD40 and CD80 (Fig. 8 A). However, surface expression of these molecules was significantly lower in cells which were subsequently stimulated with LPC (Fig. 8 A). Treatment with PC did not affect their expression (Fig. 8 A). These results suggested that stimulation of TLR-primed cells with LPC might interfere with the induction of MHC class II and co-stimulatory molecules by LPS. However, further analysis revealed that treatment with LPC also downregulated surface expression of CD11c and CD11b not only from TLR-primed cells but also from unprimed cells (Fig. 8 B, C). As ATP had been previously shown to bring about downregulation of cell surface expression of several molecules including MHC class II through protease-dependent release of vesicles (*25*–*28*), we tested if downregulation of cell surface molecules brought about by LPC was also driven by eATP and proteases. Our results showed that that indeed was the case. Treatment with apyrase or a cocktail of protease inhibitors reversed downregulation of CD11b and CD11c brought about by LPC (Fig. 8 C, D).

**Figure 8:**
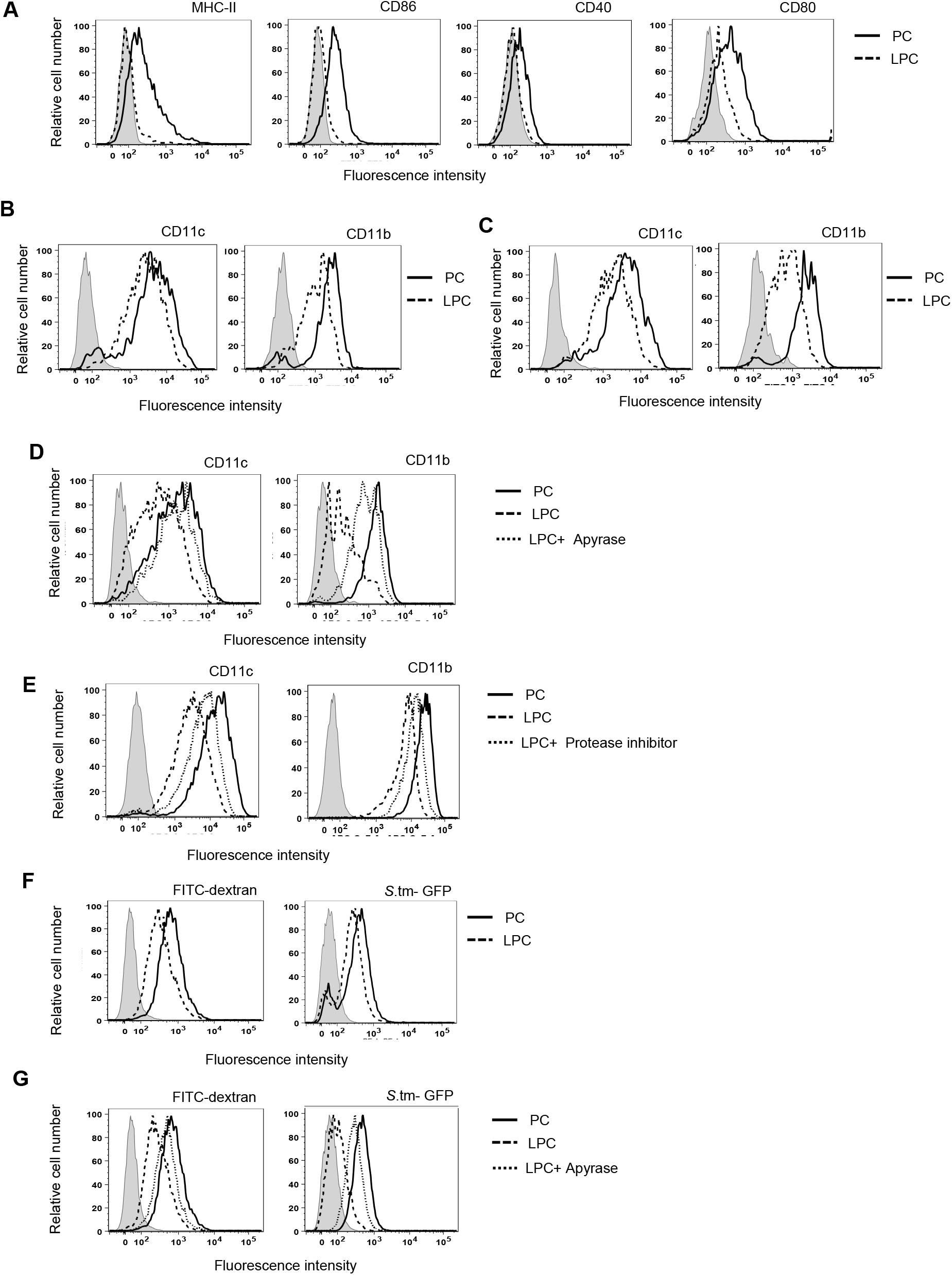
LPC-triggered eATP downregulates surface expression of key immune molecules and reduces phagocytic capability of macrophages. (**A**) Mouse peritoneal macrophages plated at a density of 5×10^5^ cells per well in RPMI-10 were stimulated with LPS (1 μg/ml) for 6 h. Cells were washed with RPMI and incubated with LPC or PC diluted in RPMI. 3 h later, cells were stained with FITC labelled antibodies against MHC-II, CD86, CD40, CD80 or isotype matched antibody. Mouse peritoneal macrophages were (**B**) left untreated or (**C**) stimulated with LPS (1 μg/ml) for 6 h. Cells were washed with RPMI and incubated with LPC or PC diluted in RPMI. 3 h later, cells were stained with APC labelled anti-CD11c, APC-Cy7 labelled anti-CD11b or isotype matched control antibody. (**D, E**) LPS-primed mouse peritoneal macrophages were washed with RPMI and pretreated with (**D**) apyrase, or (**E**) a cocktail of protease inhibitors before incubating with LPC for 3 h. Cells were washed and stained for CD11c or CD11b. (**F, G**) LPS-primed peritoneal macrophages were treated with LPC in the (**F**) absence or (**G**) presence of apyrase. Cells were washed and incubated with FITC-dextran (2 mg/ml) or GFP-labelled *Salmonella* typhimurium (25 MOI) for 1 h. Cells were analyzed by flow-cytometry (BD-Verse) after gating on live cells. Shaded histograms show isotype control (Fig. 8 A, B, C, D) or unstained cells (Fig. 8 E, F). Data are representative of 2-3 independent experiments.

LPC also reduced the ability of TLR-primed macrophages to phagocytose dextran beads or *Salmonella* (Fig. 8 E). PC did not affect phagocytosis of beads or bacteria by LPS-primed cells (Fig. 8 E). This reduction was also mediated by eATP as apyrase restored the phagocytic ability of LPC-treated cells (Fig. 8 F).

## Discussion

Chronic pathological disorders are characterized by persistent inflammation contributed significantly by DAMPs that are produced as a result of stress or cellular damage. LPC is one such DAMP whose levels have been shown to go up during many chronic diseases including atherosclerosis, multiple sclerosis and Alzheimer’s disease (*6*). LPC promotes platelet activation and vascular inflammation (*5*). It activates glial cells to produce proinflammatory cytokines (*8*, *9*). LPC has also been shown to cause demyelination *in vitro* and *in vivo* (*29*). Under normal physiological conditions, LPC engages the GPCR, G2A to perform various cellular functions (*2*). Our recent results show that serum-borne LPC can engage G2A to augment inflammatory responses from TLR – primed macrophages and epithelial cells (*30*). At higher concentrations, reminiscent of inflammation, LPC brings about caspase-1 dependent release of IL-1β from monocytes and neuronal cells through stimulation of NLRC4 and NLRP3 inflammasomes *(8)*. Our results reveal that the first critical step by which LPC initiates this caspase-1-dependent IL-1β release is the activation of release of ATP from cells. LPC triggers this release through priming of the pannexin-1 channel. However, the exact pathway by which LPC stimulates this channel is not clear at the moment. Pannexin-1 is normally activated either through proteolysis of c-terminal inhibitory region by caspases, or through a transient change in its conformation (*31*). In our analysis, ATP-release from macrophages in response to LPC seems to be independent of caspases. Therefore, LPC either engages one or more previously unrecognized proteases to produce c-terminal cleavage and activate pannexin-1, or brings about a conformational change through direct or indirect interaction with the channel. Future investigations should clarify the exact mechanism. Following ATP-release, LPC-treated cells engage one or more purinergic receptors to bring about caspase-1 activation and release of IL-1β. ATP, which is also a DAMP, normally produces caspase-1 activation from TLR-primed cells at mM concentrations but the amount of ATP that was released by LPC-stimulated cells was hundred to thousand times lower (*32*). Therefore, LPC not only triggers release of ATP from cells but it very likely also reduces the threshold at which purinergic receptors respond to this DAMP. Since peritoneal macrophages do not release IL-1β in response to ATP in the absence of P2X7R (*33*), our results suggest that it is this particular purinergic receptor that gets activated with LPC. Several mechanisms have been proposed for the activation of this receptor with ligands other than ATP (*34*). These include displacement of inhibitory cholesterol and direct activation through binding with lipid-interaction motifs in its c-terminus (*35*). LPC could activate P2X7R through either of these two mechanisms. Remarkably, ATP-release by LPC-treated cells was not only involved in bringing about caspase-1 mediated IL-1β production but it also produced caspase-1 independent cell death and promoted secretion of IL-12p70 and generation of low phagocytic and poorly co-stimulatory phenotype in a subset of macrophages. The latter phenotype would closely resemble the inflammatory macrophages with reduced capability to clear cellular debris seen in atherosclerotic plaques (*4*, *10*, *36*). Taken together, our data demonstrate an indispensable role for ATP-release in generating inflammatory responses with LPC and suggest that this ATP might contribute to cell damage and inflammation through more than one pathway. Interestingly, both LPC and eATP are considered as ‘find me signals’ that can bring about recruitment of inflammatory cells to a site of cellular damage (*37*, *38*). Our results raise the possibility that the function of LPC to serve as ‘find me signal’ might in fact be mediated through eATP.

Extracellular ATP has been implicated in the pathogenesis and promotion of a large number of chronic inflammatory disorders including inflammatory bowel disease, atherosclerosis and neurodegeneration, and more recently in hypertension associated increased immune responses (*39*), and it has been suggested that targeting purinergic signaling may be an important therapeutic option for these disorders (*40*, *41*). Our findings advocate that interfering with purinergic signaling might also be a therapeutic strategy for the management of chronic inflammatory disorders that are promoted by LPC.

## Materials and methods

### Cell lines, antibodies and other reagents

Human monocytic cell line, THP-1, was obtained from the American Type Culture Collection (ATCC, USA). Murine microglial cell line, N9, was kindly provided by Anirban Basu, National Brain Research Center, Manesar, India. *Salmonella* enterica serovar Typhimurium SL1344 (*S.* Typhimurium) was provided by Emmanuelle Charpentier, University of Vienna, Austria (now at the Max Planck Institute of Infection Biology, Berlin). GFP-expressing SL1344 was provided by Amitabha Mukhopadhyay, Cell Biology Laboratory, National Institute of Immunology, New Delhi, India. LPC and PC were obtained from Avanti Polar Lipids, U.S.A. LPS isolated from *Salmonella* typhimurium, Pam3CSK4, suramin, BAPTA-AM, EGTA, Phorbol 12-myristate 12-acetate (PMA), Pannexin-1 inhibitor (probenecid and carbenoxelone), P2XR inhibitors (A740003, KN-62, CBB-G250 and oxidized ATP), ATP, apyrase and FITC-dextran were obtained from Sigma-Aldrich Co., U.S.A. Flagellin was prepared from *Salmonella* Typhi as described previously (*42*). ELISA kits for detection of mouse IL-1β was from BioLegend and for mouse IL-12p70 and human IL-1β were from BD. Pertussis toxin, caspase-1 inhibitors z-YVAD-fmk, caspase-3 inhibitor and pan-caspase inhibitors were obtained from Calbiochem, U.S.A. ATP bioluminescense assay kit HS-II and Protease inhibitor cocktail was from Roche. Flou-4AM was from Thermo Fisher Scientific. Non-radioactive cytotoxicity kit for determination of cell death was obtained from Promega Corporation, Wisconsin, U.S.A. Fluorophore-conjugated antibodies against mouse MHC-II, CD-80, CD-86, CD40, CD11c, CD11b and isotype matched control antibodies were obtained from e-biosciences, USA.

### Mice

C57BL/6 inbred mice were obtained from Jackson Laboratories, U.S.A, and maintained in the Small Animal Facility of the National Institute of Immunology. Experiments with mice were carried out in accordance with the guidelines provided by the Institutional Animal Ethics Committee of NII.

### Isolation of mouse cells

6-8-week-old C57BL/6 mice were injected with 4% thioglycollate and 3 days later, cells were taken out of the peritoneum using ice-cold PBS. Cells were subjected to plastic adherence in cell culture dishes for 4 hours. Non - adherent cells were removed and adherent cells were used as macrophages throughout the study unless otherwise mentioned.

### Activation of cells with LPC, and determination of cytokines and LDH

Cells were primed with LPS or Pam3CSK4 (1μg/ml) in duplicate in a 24 well tissue culture plate at a density of 5×10^5^ cells per well for 6 h at 37°C (in a humidified atmosphere with 5% CO_2_). Cells were washed extensively with serum - free RPMI-1640 and stimulated with LPC or PC for 3 h. In some experiments, TLR-primed cells were treated with various inhibitors for 30 minutes before incubating with LPC for another 3 h. The levels of IL-1β and IL-12p70 were determined in triplicate in cell culture supernatants by ELISA. The levels of LDH were determined by LDH cytotoxicity assay kit.

### Determination of ATP-release

Cells seeded at a density of 5×10^5^ cells per well were washed free of serum and stimulated with LPC or PC (50 μM). Supernatants were collected at different time points, boiled at 90°C for 2 minutes and ATP was quantified using ATP bioluminescence assay kit HS-II in accordance with the manufacturer’s instructions.

### Determination of intracellular calcium

Peritoneal macrophages seeded at a density of 2×10^5^ cells per well were washed free of serum and loaded with calcium indicator dye FLOU-4AM (5 μM dissolved in Ca^2+^free HBSS) for 30 minutes at 37°C in dark in a humidified atmosphere with 5% CO_2_. Cells were washed and rested for another 30 min. CaCl2 (2 mM) was added and cells were stimulated with LPC or PC (50 μM). Calcium flux was recorded after 5 minutes. Images were taken under a fluorescence microscope (Nikon TE-2000) and imported into adobe photoshop. In some experiments, calcium flux was monitored in presence of apyrase.

### Analysis of *in vivo* response with LPC

Mice were injected intraperitoneally with LPS (7 μg / mouse) and 10 h later, LPC (200 μg / mouse) was administered intraperitoneally. Peritoneal lavages and sera were collected after 2 h and 4h respectively. Cell-free peritoneal lavages and sera were analyzed for IL-1β by ELISA. In some experiments, mice were given apyrase (15 U) along with LPC.

### Microscopic examination of cells

TLR-primed cells seeded at a density of 5×10^5^ cells per well were washed free of serum and stimulated with LPC or PC in presence or absence of different inhibitors for 3 h. Cells were visualized under a fluorescence microscope (Nikon TE-2000) and images were acquired using ACT software and imported into adobe photoshop.

### Flow cytometric analysis of surface expression of molecules and phagocytosis

TLR-primed cells were incubated with LPC or PC in presence or absence of apyrase for 3 h. Cells were harvested and incubated with fluorophore-conjugated antibodies against mouse MHC-II, CD-86, CD40, CD80 CD11c, CD11b or isotype-matched control antibodies for 1 h at 4°C (BD biosciences, San Jose, CA) or and incubated with FITC-dextran or non-invasive GFP-expressing *Salmonella* for 1 h at 37°C. Cells were washed in PBS and analyzed by flow cytometry (FACSVerse®, Becton Dickinson Immunocytometry Systems, San Jose, CA). Data was plotted using Flowjo software.

### Statistical analysis

Student’s *t*-test was used to calculate p-values by 2 tails with type 3 test for unequal variances. P value of less than 0.05 was considered statistically significant. Data are expressed as mean ± SD. Error bars represent standard deviation.

## SUPPLEMENTARY MATERIALS

**Fig. S1.** Suramin and not pertussis toxin inhibits IL-1β from TLR-primed and THP-1 and N9 cells stimulated with LPC.

**Fig. S2.** Apyrase inhibits LPC-induced IL-1β from THP-1 and N9 cells.

**Fig. S3.** LPC-induced IL-1β depends on calcium mobilization, Syk, JNK and potassium efflux.

**Fig. S4.** LPC-induced extracellular ATP brings about calcium mobilization.

**Fig. S5.** LPC brings about YO-PRO1 uptake inhibitable with apyrase.

**Fig. S6.** Inhibitors of purinergic signaling abrogate LPC-induced IL-1β from THP-1 and N9 cells.

## Acknowledgements

We thank Drs Rahul Pal, Devinder Sehgal, Pushkar Sharma and Mohammed Asif for their valuable suggestions in the course of this study, and members of the Ayub laboratory for insightful discussions. This work was funded through the National Institute of Immunology by the Department of Biotechnology, Government of India.

## Author contributions

A.Q. conceived and supervised the study. S.I. and A.Q. designed experiments. S.I. performed experiments and prepared data for publication. S.I. and A.Q. analyzed the data, and wrote the manuscript.

## Conflict of interest

The authors declare no financial or commercial conflict of interest.

## Notes

### Competing Interest Statement

The authors have declared no competing interest.

