## Supplementary material for "ATP-release drives inflammation with lysophosphatidylcholine"

**Running title:** LPC triggers release of ATP for inflammation

**Key words:** Lysophosphatidylcholine, ATP, caspase-1, pannexin-1, purinergic receptor

**Supplementary Fig 1: Suramin and not pertussis toxin inhibits IL-1β from TLR-primed THP-1 and N9 cells stimulated with LPC.**

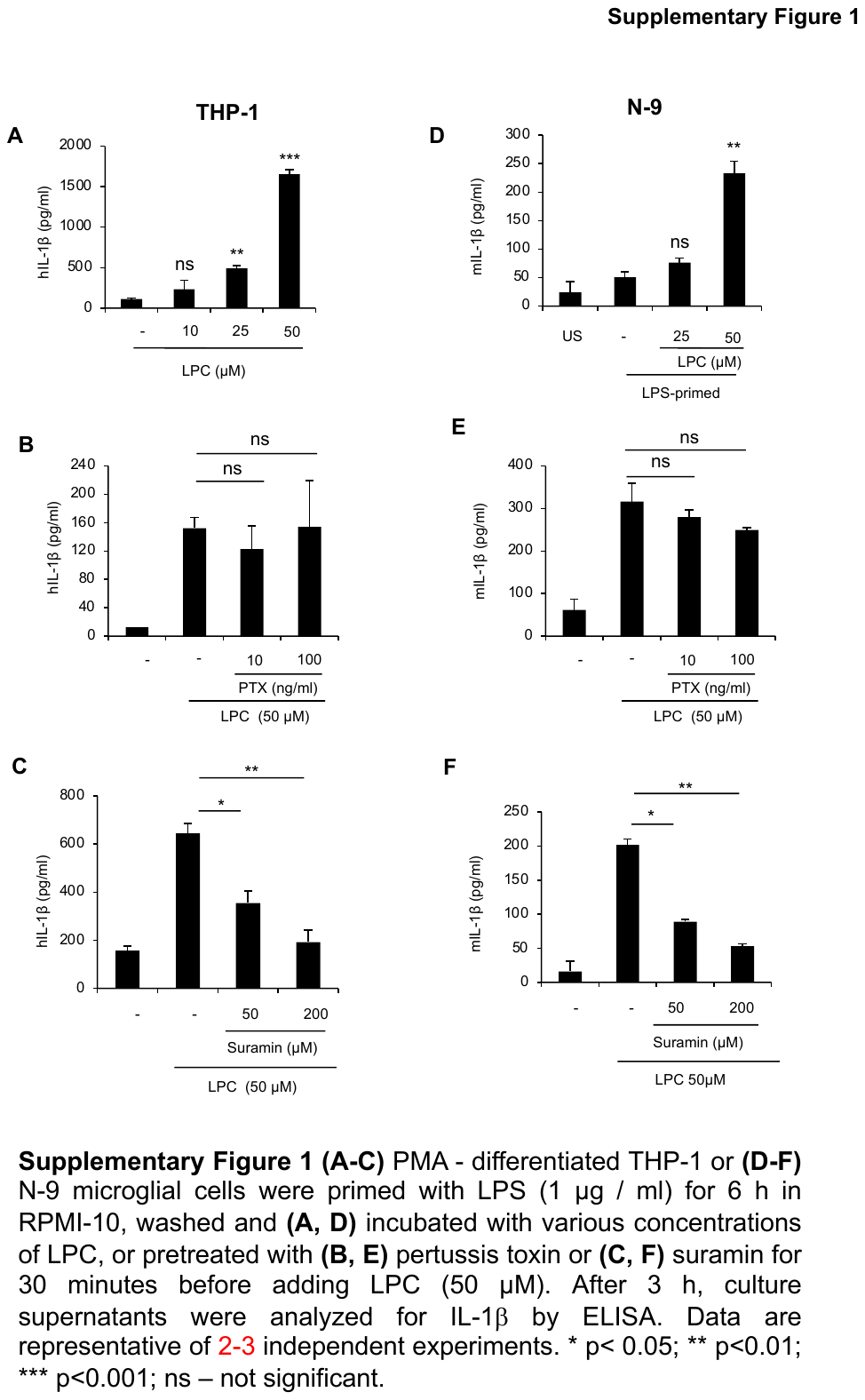

(**A-C**) PMA - differentiated THP-1 or (**D-F**) N-9 microglial cells were primed with LPS (1 μg / ml) for 6 h in RPMI-10, washed and (**A, D**) incubated with various concentrations of LPC, or pretreated with (**B, E**) pertussis toxin or (**C, F**) suramin for 30 minutes before stimulating with LPC (50 μM). After 3 h, culture supernatants were analyzed for IL-1β by ELISA. Data are representative of 2-3 independent experiments. * p< 0.05; ** p<0.01; *** p<0.001; ns – not significant.

**Supplementary Fig 2: Apyrase inhibits LPC-induced IL-1β from THP-1 and N9 cells**

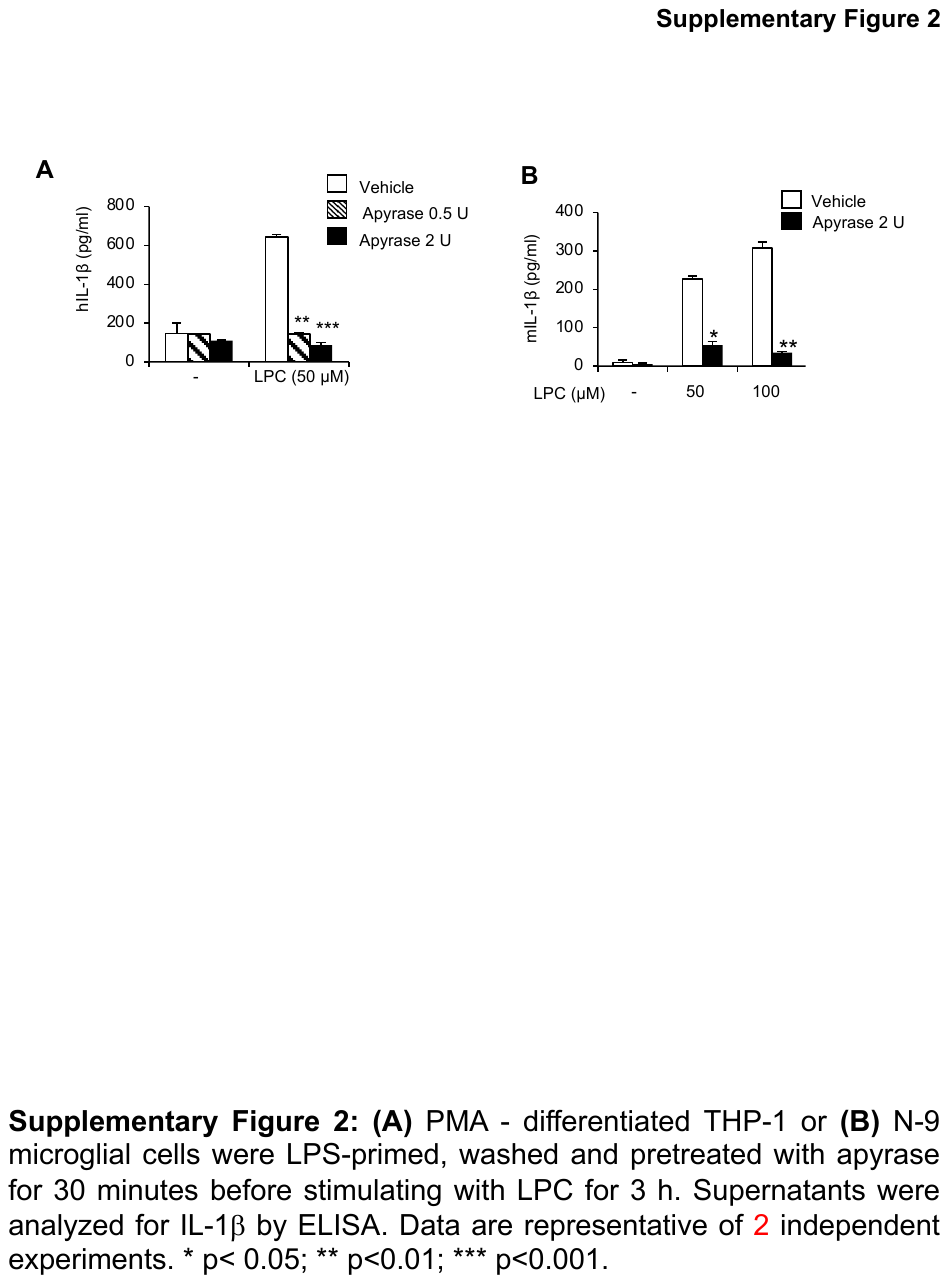

(**A**) PMA - differentiated THP-1 or (**B**) N-9 microglial cells were LPS-primed, washed and pretreated with apyrase for 30 minutes before stimulating with LPC for 3 h. Supernatants were analyzed for IL-1β by ELISA. Data are representative of 2 independent experiments. * p< 0.05; ** p<0.01; *** p<0.001.

**Supplementary Fig 3: LPC-induced IL-1β depends on calcium mobilization, Syk, JNK and potassium efflux.**

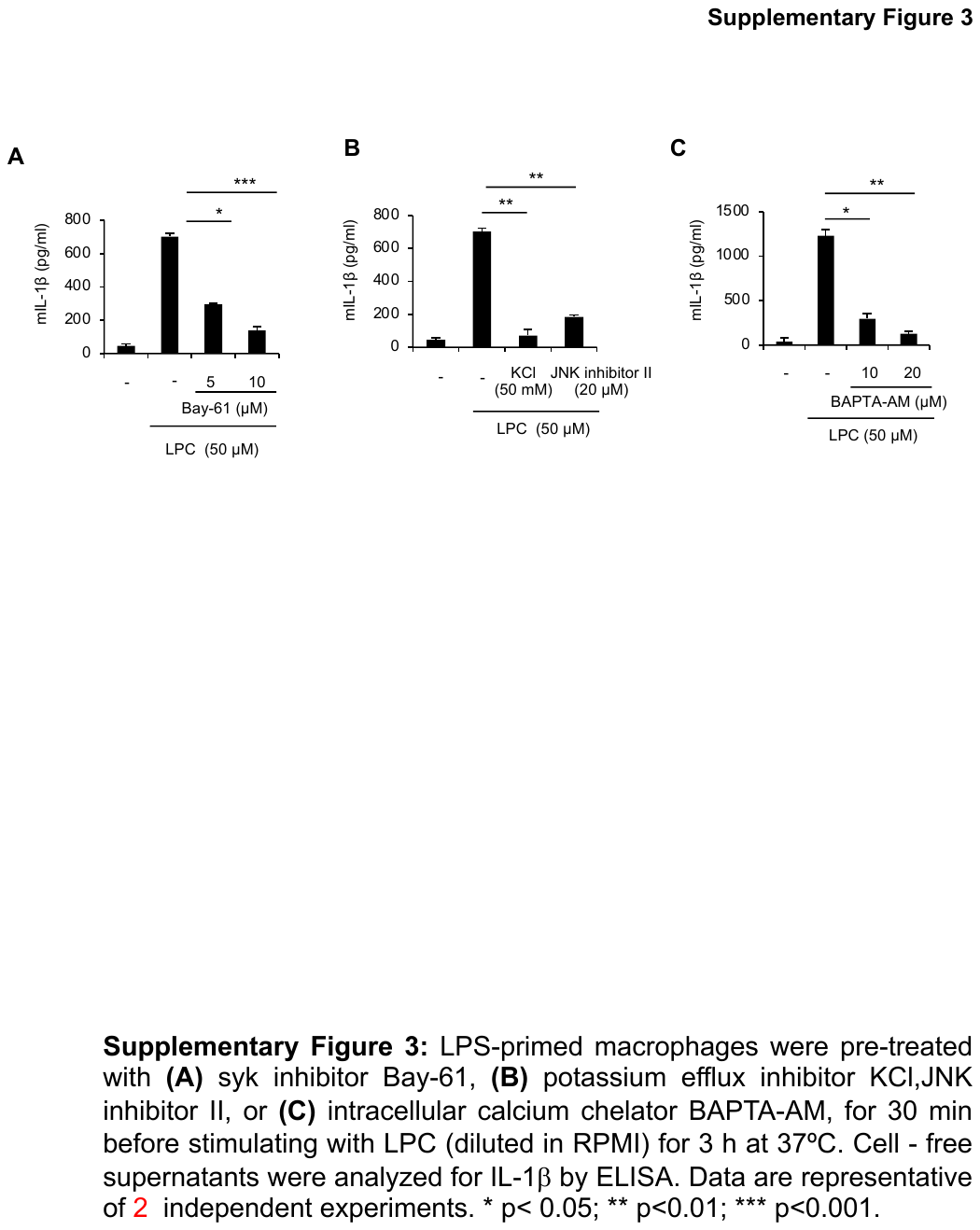

LPS-primed macrophages were pre-treated with (**A**) syk inhibitor Bay-61, (**B**) potassium efflux inhibitor KCl, JNK inhibitor II, or (**C**) intracellular calcium chelator BAPTA-AM, for 30 min before stimulating with LPC (diluted in RPMI) for 3 h at 37ºC. Cell - free supernatants were analyzed for IL-1β by ELISA. Data are representative of 2 independent experiments. * p< 0.05; ** p<0.01; *** p<0.001.

**Supplementary Fig 4: LPC-induced extracellular ATP brings about calcium mobilization.**

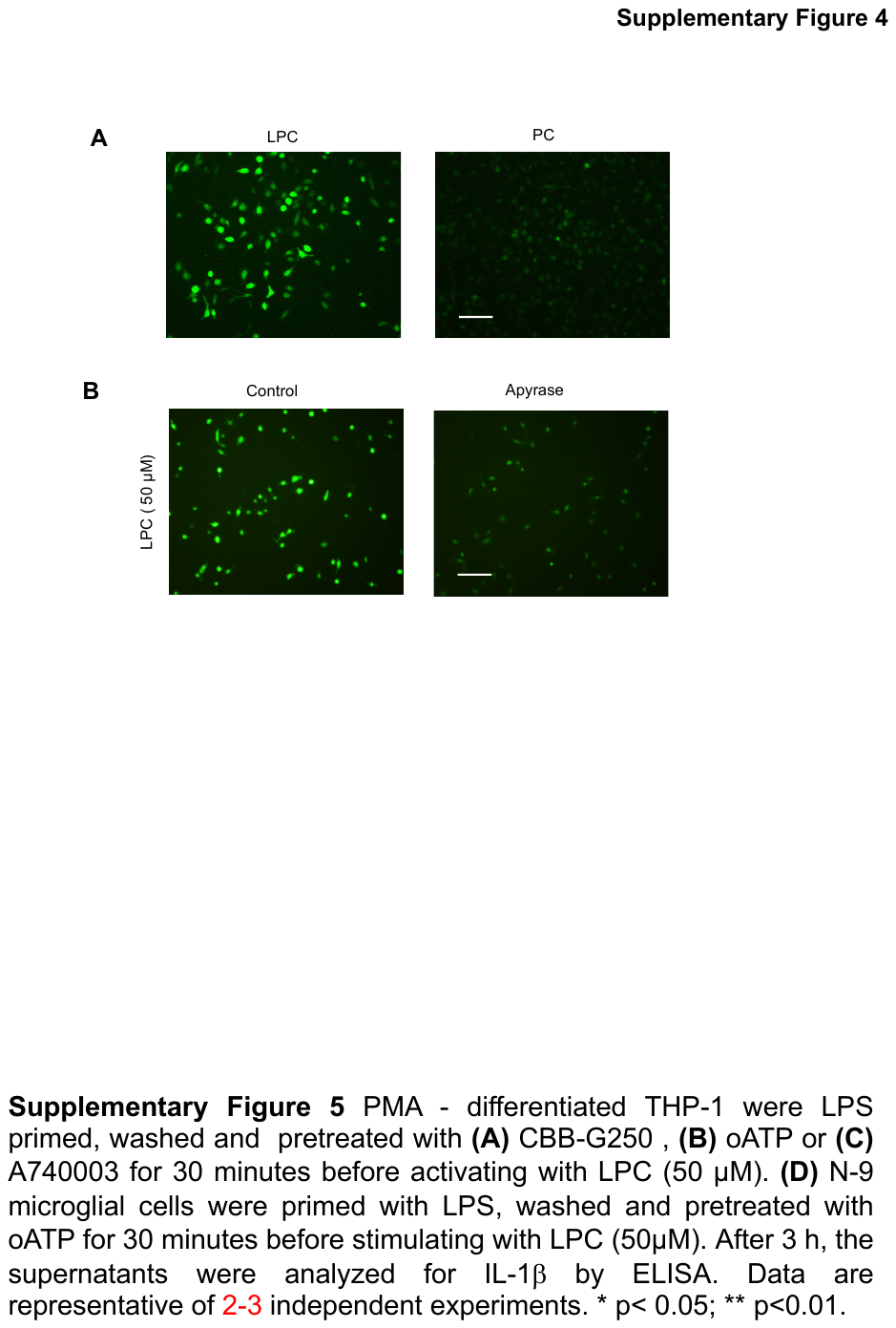

(**A, B**) Murine macrophages were loaded with calcium indicator dye Fluo-4AM in HBSS for 30 min. Cells were washed and allowed to rest for 30 min in HBSS. CaCl_2_ was added before stimulating cells with LPC or PC (50 μM) in the absence or presence of apyrase. Images were captured after 5 min in a fluorescence microscope (Nikon TE-2000) using ACT software and imported into adobe photoshop. Scale bar- 50 μm. Data are representative of 2 independent experiments.

**Supplementary Fig 5: LPC brings about YO-PRO1 uptake inhibitable with apyrase.**

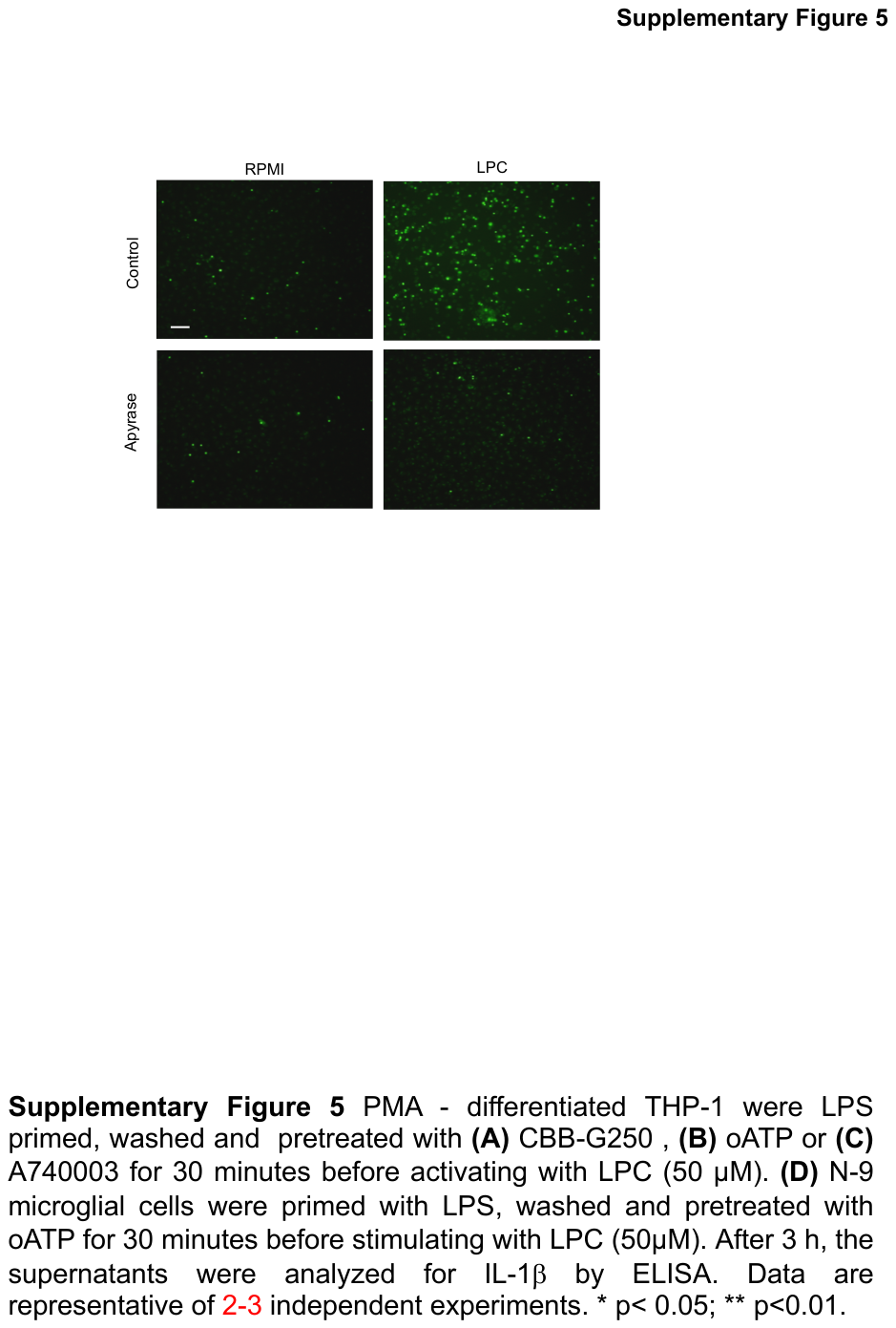

Murine macrophages were washed free of serum and treated with apyrase for 30 minutes before stimulating with LPC (50 μM). 3 h later, YO-PRO1 (1 μM) was added to cultures and images were taken in a fluorescence microscope (Nikon TE-2000) using ACT software and imported into adobe photoshop. Scale bar- 100 µm. Data are representative of 3 independent experiments.

**Supplementary Fig 6: Inhibition of purinergic signaling abrogates LPC-induced IL-1β from THP-1 and N9 cells.**

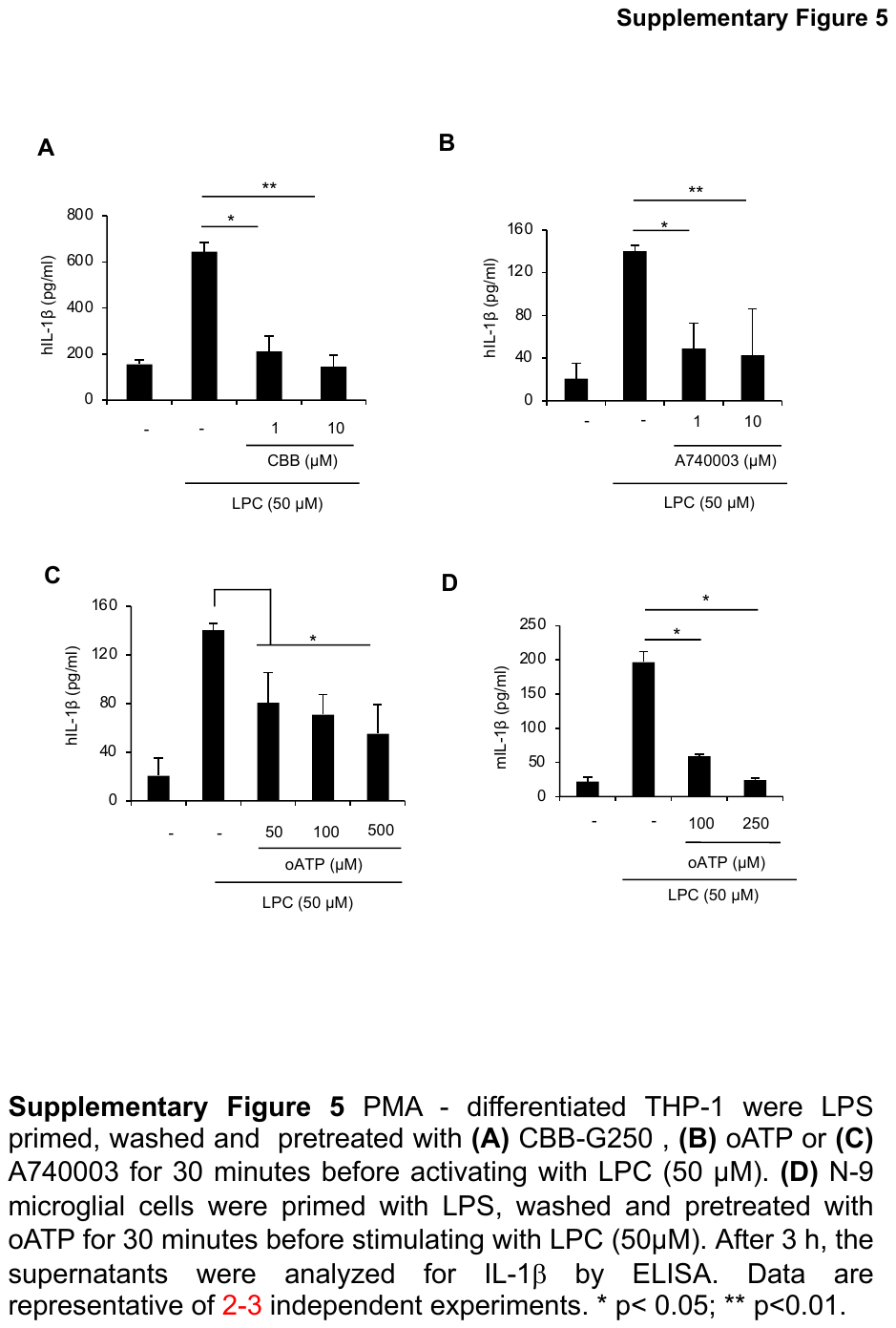

PMA - differentiated THP-1 were primed with LPS, washed and pretreated with (**A**) CBB-G250, or (**B**) A740003 for 30 minutes or with (**C**) oATP for 2 h before activating with LPC (50 μM). (**D**) N-9 microglial cells were primed with LPS, washed and pretreated with oATP for 2 h before stimulating with LPC (50 μM). After 3 h, the supernatants were analyzed for IL-1β by ELISA. Data are representative of 2-3 independent experiments. * p< 0.05; ** p<0.01.
